# Abiotic environmental adaptation in vertebrates is characterized by functional genomic constraint

**DOI:** 10.1101/726240

**Authors:** Katharina C. Wollenberg Valero, Joan Garcia-Porta, Iker Irisarri, Lauric Feugere, Adam Bates, Sebastian Kirchhof, Olga Jovanović Glavaš, Panayiotis Pafilis, Sabrina F. Samuel, Johannes Müller, Miguel Vences, Alexander P. Turner, Pedro Beltran-Alvarez, Kenneth B. Storey

## Abstract

Understanding the genomic basis of adaptation to different abiotic environments is important for understanding organismal responses to current short-term environmental fluctuations. Using functional and comparative genomics approaches, we here investigated whether genomic adaptation to a set of environmental parameters is contingent across vertebrate genomes or, alternatively, contains an element of evolutionary constraint that would be evident through recurrent involvement of specific subsets of genes and functions in adaptation to similar environments. We first identified 200 genes with signatures of selection from transcriptomes of 24 species of lacertid lizards with known adaptations in preferred temperature, correlated with thermal environment experienced by these lizards in their range. In order to discern genes adapting to climate from other selective factors, we then performed a meta-analysis of 1100 genes with signatures of selection obtained from-omics studies in vertebrate species adapted to different abiotic environments. We found that this gene set formed a tightly connected interactome which was to 23% enriched in predicted functions of adaptation to climate and to 18% involved in organismal stress response. We found a much higher degree of recurrent use of identical genes (43.6%) and functional similarity than expected by chance, and no clear division between genes used in ectotherm and endotherm physiological strategies. 171 out of 200 genes of Lacertidae were part of this network, indicating that a comparative genomic approach can help to disentangle genes functionally related to adaptation to different abiotic environments from other selective factors. These results furthermore highlight an important role of genomic constraint in adaptation to the abiotic environment, and narrows the set of candidate markers to be used in future research on environmental adaptability related to climate change.

**Significance Statement / Nontechnical summary:** While the convergent evolution of phenotypes in similar environments is a well-studied phenomenon, the genomic basis of such common phenotypes and physiologies is still enigmatic. The prevalent notion is that re-use of the same genes to adapt to similar environments in different species is about as likely as winning the lottery – but organismal systems are also, to some extent, comparable between different species such as man and fruit fly through shared genes and gene functions. In this paper, we test whether constraint or contingency is more prevalent in genomic adaptation of vertebrates to aspects of their abiotic environment. We find evidence for strong functional constraint and stress responsiveness of the genes involved, which might help understand how currently experienced stress under changing climates may result in future adaptation.

## Introduction

The sheer number of genes and alleles (1) within a genome, the fact that many traits are polygenic (2) and that genomes can evolve via alternative pathways (3) theoretically provide infinite possibilities for adaptations to evolve. As a result of this, genomes are expected to reflect a high degree of contingency (evolutionary unpredictability (4, 5)). However, many studies have noted that there are less realized adaptations than expected, which means that there could be constraints at the genome level that could lead to evolutionary determinism (more predictable outcomes (4)) (1). One such element of constraint could be explained through the inertia generated by the interaction of genes that perform a common function upon which selection acts (gene functional constraint) (6–8). In this paper, we address the question of how prevalent such constraint is at the genomic level, on the example of adaptation to aspects of the abiotic environment.

First, constraint in adaptation to a common selective pressure (e.g., the abiotic environment) could be detected by comparing the similarity of gene functions involved in adaptation across different taxa. Adaptation to different or novel abiotic environments involves changes in traits which, in similar environments, often converge (9). Traits of the phenotype or physiology are based on groups of gene products operating within defined functions (7). Through common descent, this relationship between specific genes, their functions, and traits may be conserved across different lineages. A high degree of conservation can constrain the number of genes performing a function. As a result, the number of genes that may be modified by selection on that function to adapt across these different lineages may be constrained (Fig. 1) (10). For example, adaptation to climate in vertebrate ectotherms such as squamate reptiles, is directed through a specific set of gene functions with evidence that similar genes are involved (8, 11, 12). In vertebrate ectotherms, these functions contain many genes involved in the conserved stress response (13), e.g., heat shock proteins (HSPs) (14). Other candidates for environmental adaptation are genes active within a cancer microenvironment which likewise represents aspects of a stressful environment such as hypoxia and acidic pH. Furthermore, cellular stress such as oxidative stress often ends in apoptosis (15) and consequently, genes related to the cellular pathways of apoptosis, stress and inflammation (dubbed “Zombie genes” (16)) could also be candidates for environmental adaptation. Functionally similar genes can also be recognized through similar epiproteomic signals, such as protein post-translational modifications (PTMs (17)).

**Figure 1.**
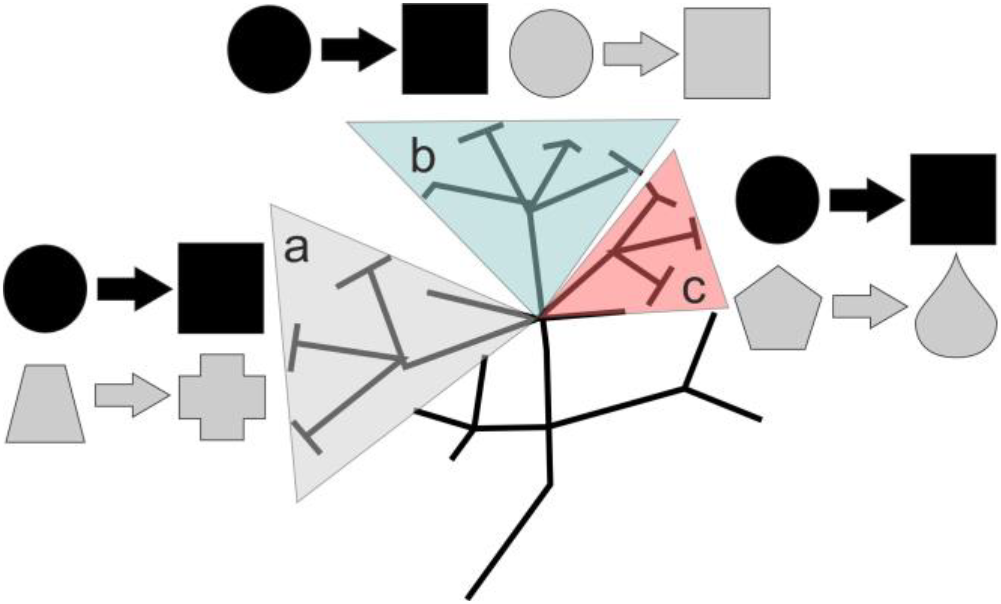
Contingency versus gene functional constraint at the gene product level. Organisms use gene products (shapes) to perform functions (arrows) at the molecular level. **Expectation of constraint:** Since they share a common ancestor, it is likely that these molecular functional systems likewise share common ancestry. If a specific function (such as temperature homeostasis) is under strong balancing selection, it is unlikely that over time genes with this function get exapted to other functions, or that new genes get recruited into this function (shown here as black symbols). In different lineages (a-c), the molecular function will therefore be performed by the same gene products. If adaptation then occurs due to a novel environment (e.g., a different environmental temperature), it is likely that the adaptation (in the form of changed gene sequence or expression level) will be constrained to the set of genes which has evolved to perform that function. If several lineages are compared, we expect to find genes being recruited multiple times to adapt to this change in thermal environment, i.e., recurrently. **Expectation of contingency:** In contrast, if the function of temperature homeostasis is under relaxed constraint (shown as grey gene symbols), the system will be modified through evolution to recruit and lose different gene products. A novel abiotic environment will lead to adaptation in different EAGs in different lineages. Consequently, the degree of genes recurrently involved in adaptation will be very low and more comparable to the likelihood of drawing genes from the genome by chance.

Second, evidence for constraint can be estimated through the number of identical genes that are involved in the functions related to specific aspects of adaptation. Under gene functional constraint, we expect that genes cannot easily be recruited or lost from their function, and thus we would expect to find adaptations to similar selective pressure to be concentrated in a subset of genes performing these functions across different organisms. Under strong constraint, or with just a small number of genes performing these relevant functions, this could even lead to the re-use of the same genes in adaptation to similar environments. We here call this process recurrence, or recurrent evolution in cases where the same genes are recruited or modified for adaptation in non-sister lineages. One of the best-known examples of recurrent evolution based on modification of identical genes in different species is altitude adaptation in the Himalayas. High-altitude yak, Tibetan humans and their dogs all show adaptations in the ADAM17 gene related to the physiological function of hypoxia tolerance (18, 19) compared to their low-altitude relatives. An example for recurrent evolution independent of common descent are the antifreeze glycoprotein genes in Antarctic notothenioid fish and Arctic cod. Antarctic fishes re-purposed the glycoprotein sequence from a different ancestral function (20), whereas the same functional gene sequence in Arctic cod evolved *de novo* from a non-coding genomic region (21). Alternatively, through modification, different genes could perform these functions in different organisms, which would indicate the constraint on genes contributing to adaptation is more relaxed. Complete relaxation of the gene functional constraint would mean that any gene could perform any function in different lineages and genes adapting to different environments across lineages will be different across lineages, and will have less functional similarity (Fig. 1).

Third, constraint can be identified through comparing organisms with obvious differences in physiological strategy. Strong constraint will keep gene function and gene identity similar even across divergent physiologies and evolutionary lineages, whereas relaxed constraint will lead to more pronounced genetic differences across these groups. Especially in those vertebrates where development in the majority of taxa occurs outside the parental body, climate can be a strong selective force for both cold (12, 22) and hot (23) climate stress and -adaptation. While endothermic vertebrates (including humans) have been well studied from an -omics perspective (24), the genomic basis of environmental adaptation in vertebrate ectotherms is generally understudied (25), with only a few studies incorporating genome-wide scans for environmental adaptation-relevant genes to date (12, 26–28).

The first aim of this contribution is therefore to better understand the genomic basis of environmental adaptation in vertebrate ectotherms in one such group, squamate reptiles belonging to the Western Palearctic lacertid lizards (family Lacertidae). We use an RNA sequencing data set we previously generated for phylogenomic analysis of this group (29) to newly identify genes with signatures of selection across 24 species representing the lacertid diversity. Informed by the physiological adaptations in preferred body temperature (T_pref_) with bioclimate identified by us in a previous study (29), we hypothesize that at least some of these genes will harbor genomic adaptations to cold across the climatic gradient they inhabit (30).

To identify the extent of constraint in adaptation to the abiotic environment across vertebrates, we then compare the functions of genes putatively involved in environmental adaptation (in the following called environmental adaptation genes, EAG) across vertebrates, mined from 25 -omics studies of adaptation and 75 -omics studies of stress. Under the premise of gene functional constraint, we expect that EAGs identified in different lineages are tightly functionally connected, fall within predefined functions for climate adaptation (11, 12, 31) and are expressed or modified in cellular functions related to stressful conditions (here: environmental stress, apoptosis, cancer). We also expect functional similarity of both lacertid and other vertebrate EAGs to be higher than expected by chance. Lastly, we aim to understand the extent of genomic recurrence across endotherm and ectotherm vertebrates including Lacertidae. We quantify the genes that are identified in more than one species, and compare their frequency of occurrence against a random draw simulation representing the absence of constraint as null hypothesis. Here, we expect to find both functional similarity as well as evidence for constraint in the modification of EAGs across different taxa.

## Results

Among 5,498 ortholog genes compiled from the transcriptomes of 24 lacertid taxa (29), we identified 200 genes to have evolved under episodic positive diversifying selection. The estimated median ω (dN/dS ratio) per branch for these 200 genes was plotted onto a maximum likelihood phylogeny (Fig. 2a). Branches splitting from basal nodes, except *Podarcis*, showed the highest median values of ω. The majority of 10>ω>1 genes were located on the (long) terminal branches. In contrast, ω ≥ 10 genes were concentrated both along the phylogeny backbone and on a set of terminal branches. Based on previous literature meta-analyses and experimental confirmation of genes adapting to climate, a set of candidate functions related to climate adaptation was defined (11, 12, 31). We tested whether newly identified genes were also functionally enriched in these functions. The Lacertid genes were significantly enriched (Fig. 2b) in several previously defined candidate functions potentially related to climate adaptation (11, 12, 31), and in cytoskeletal processes as additional, not predicted functional category (Tables S1-S2).

**Figure 2.**
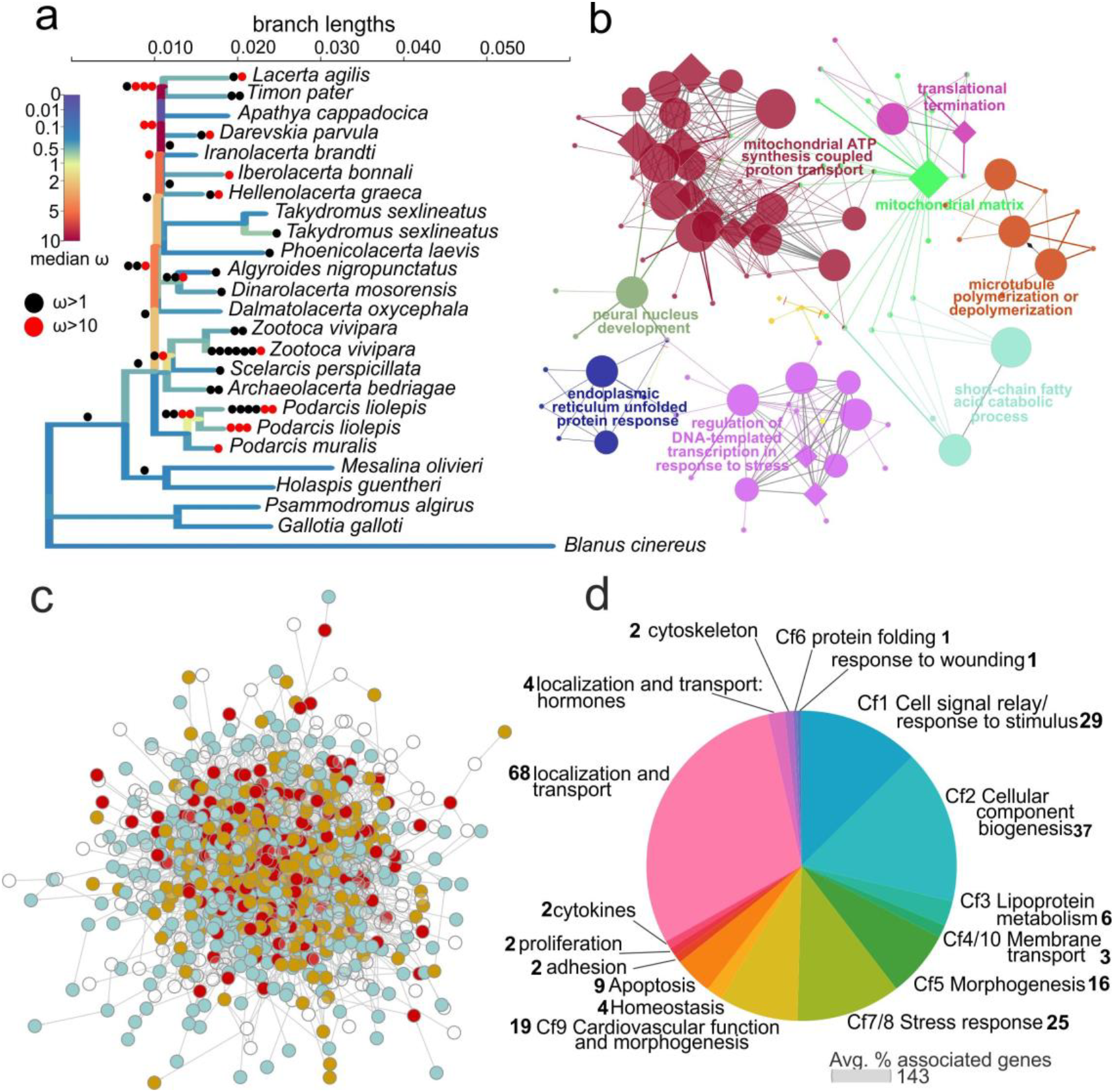
Phylogenetic and functional properties of lacertid and vertebrate EAGs. (a) Visualization of median selection coefficient (ω) on RNAseq maximum likelihood phylogram of 24 lacertid species for 200 genes under episodic positive diversifying selection. Genes with 10>ω>1 and ω>10 are shown along the branches as circles. (b) Function network of significantly enriched functions for these genes in Lacertidae (circles - biological process; diamond - molecular function). (c) Functional genomic network for environmental adaptation generated from 902 genes across endotherm and ectotherm vertebrates (blue –genes that recurrently adapted to environmental parameters in different species; ochre –genes that are stress responsive, maroon –genes that evolved both recurrently and are stress responsive). (d) Graph of functional categories of significantly overrepresented ClueGo Gene Ontology groups, for the 902 functionally connected environmental adaptation genes, categorized by candidate functions (Cf) and additional functions. Numbers in labels refer to numbers of Gene Ontology terms within each category.

A total of 1100 putative EAGs across ectotherm and endotherm vertebrates (12 ectotherm and 17 endotherm species) were compiled from -omics studies of adaptations to different or novel abiotic environments. These genes resulted from statistical comparisons among sister clades in phylogenies, among populations, or across generational timepoints. Including the 200 Lacertidae genes, the set of 1100 genes were involved in adaptation to different environmental conditions comprised of thermal differences in latitude, altitude or season, sulfidic water in streams, osmotic differences, desiccation, hypoxia, or wet-dry environmental gradients (Tables S3). Of the 1100 putative EAGs, 902 or 83% were part of a large, tightly connected protein-protein interaction (PPI) network (Fig. 2c). Among these, 480 genes or 43.6% aligned to our definition of recurrent evolution as they were involved in environmental adaptation in multiple, not closely related vertebrate clades (Fig. 2c, Table S4). The Gene Ontology (GO) terms for this network covered all except one *a priori* predicted function potentially related to climate adaptation, which was muscle use and -development (Table S1, S5). Additional significantly enriched GO terms within this network were related to the cytoskeleton, apoptosis, and localization and transport (Table S1, S5, Fig. 2d). The latter function encompassed 68 related GO terms and 459 genes (45%). 86% or 171 of the 200 selected genes in Lacertidae were also part of this functional network and 14 of these adapted recurrently with other taxa (Fig. S1, Table S1). As outlined above, we consider it likely that at least these 902 genes (171 of Lacertidae) connected to each other by climate adaptation-relevant functions and evidence for recurrence represent true EAGs, while this is less likely for the 29 Lacertidae genes not connected to this network. To investigate the extent of constraint through gene re-use in environmental adaptation, we tested whether the number of 480 recurrently evolving genes out of 1,843 genes across all data sets (some of the 1100 individual genes were identified in more than two species) was higher than expected by chance. In 100,000 simulations, we found that in order to obtain an identical number of recurrent genes solely by chance, on average 4736.7 (± 93.4) genes would have to be drawn from a hypothetical genome of 20,000 genes. This value is significantly higher than the empirical value obtained here (Fig. 3a, Z= −30.98, p ≤ 0.00001).

**Figure 3.**
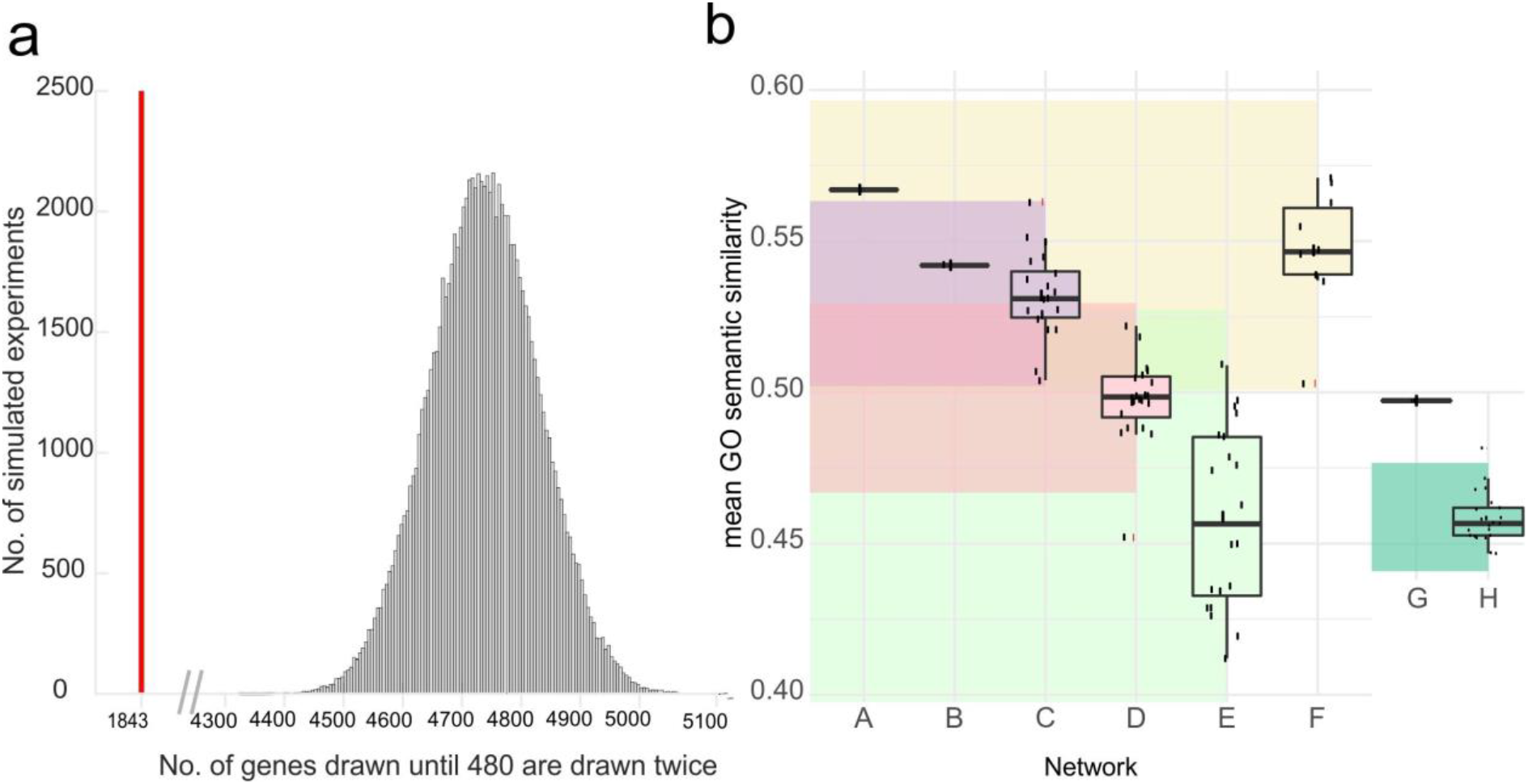
Comparison of lacertid genes under selection and vertebrate recurrently evolving EAG to randomized and simulated data. (a) Number of genes that was drawn in this study to find 480 recurrent genes (red bar) is significantly lower than the number of genes obtained in N=100,000 simulations of random draws from a hypothetical genome of 20,000 genes to yield 480 recurrent genes, as expected under no constraint (histogram, ***). **(b)** Comparisons of functional (semantic) similarity of GO terms between genes under selection in Lacertidae and random networks (A-F), and between 902 vertebrate EAG and 20 sets of 902 randomized genes (G, H). Boxes show medians, 25-75 quartiles, outliers, and jittered raw data. Transparent bands indicate modelled ±95% prediction intervals. The plot shows that the 200 genes under selection (A) have the highest functional similarity. Their semantic similarity is higher than that of 29 genes under selection not part of the network (B), of 20 random replicates of each 200 genes not under selection (C,**); of 20 random replicates of 200 genes drawn from 5,498 annotated ortholog genes with incomplete taxon sampling obtained from RNAseq (D, ***), of 20 replicates of 200 random human genes (E, ***), but not higher than 10 replicates of 451 human housekeeping genes (F); 902 vertebrate EAGs part of a functional genomic network (G) have significantly higher functional similarity than 20 random samples of 902 genes (H, ***). p values are denoted as * ≤ 0.05; ** ≤ 0.01; *** ≤ 0.001.

We then tested whether the functions and functional connectedness of vertebrate EAGs were more similar than expected by chance or properties of the sequenced transcriptomes. The vertebrate EAG network had significantly more interactions between the genes than expected if they were randomly drawn from the genome, with a PPI enrichment p-value of 2e^−13^. It also had significantly higher functional similarity, measured via the semantic similarity of its GO terms than random networks of comparable size (Fig. 3b, Z = 4.57, p = 0.00001). When likely lacertid EAGs were separately subjected to GO enrichment analysis, two additional candidate functions for climate adaptation were recovered (cell signalling and muscle-related processes; Table S1). For the 29 Lacertidae genes under selection that were not part of the network (and less likely constituting EAGs), no functional GO term enrichment could be detected (Table S1). Lacertid EAGs also had higher semantic similarity of GO terms than those 29 not part of the network, and higher semantic similarity than randomized replicates of all other comparison groups (29 genes not under selection Z = 2.5, p = 0.012, all other annotated orthologs in transcriptomes Z = 5, p ≤ 0.00001, randomly drawn genes Z = 3.6, p = 0.0003, Fig. 3b), but not compared to human “housekeeping” genes. (Fig. 3b, Z = 1, p = 0.159).

We subsequently tested whether putative EAGs, and those with evidence for recurrent adaptation, could be grouped by a priori defined candidate functions for climate adaptation, or related to stress (evidence for stress responsiveness, cancer and apoptosis), as well as PTMs. Evidence for human gene PTMs was present in 19% of EAGs, and 21% genes furthermore had the function “gene regulation - protein modification” (Table S6). EAGs overall had significantly more PTMs than randomly drawn genes (Fig. S2, Mann-Whitney W = 19572, p ≤ 2.2e^−16^), and significantly less PTMs than randomly drawn human housekeeping genes (W = 8599, p = 2.57e^−15^). These results remain significant after comparison with a randomized distribution of W (EAG to random genes Z = 443.726, p ≤ 0.001; EAG to housekeeping genes Z = 192.44, p ≤ 0.001, Fig. S2). 198 EAGs or 18% had “stress response” as a significant GO term (Table S4). Furthermore, we found empirical evidence in the literature for 329 (29.9%) genes being differentially expressed in response to abiotic environmental stress (including cold stress, heat stress, hypoxia stress, and others including hibernation, desiccation and detoxification, Table S6, the complete data is provided in Table S7). Of these, 125 (38.0%) were upregulated and 74 (22.5%) were downregulated (Fig. S3). The top three candidate functions for climate adaptation represented by these genes were “cellular component biogenesis”, “morphogenesis”, and “response to oxidative stress / stress response (incl. thermal stress)” (Table S8-S9). All of these genes were part of the functional network. Several of them were heat shock proteins; however, the two most recorded genes responding to all analyzed aspects of environmental stress across different species were the less-studied BAG3 and AHSG. AHSG furthermore responded to all four groups of stressors, together with PKLR and SQRDL (Fig. S4a). With regards to cellular pathways active in cancer as possible EAG candidate function, we found that recurrent EAGs have lower expression values in both cancer and healthy tissues (Fig. S4b), whilst ratios of gene expression between cancer and reference conditions did not differ (Table S10). Five genes of the data set were, however, involved in more than 80 cancer types. (Fig. S5, Table S11). To visually align EAGs to candidate functions, six functional dimensions of EAGs were extracted by Multiple Correspondence Analysis. Fig. 4 shows the dendrogram obtained by *pvclust* following 100,000 bootstrap replicates. Over ten hierarchical functional clusters were supported with bootstrap values >95, with functional dimensions not clearly partitioning genes into clusters that could be identified as clearly ectotherm or endotherm (for more details see Supplementary Results).

**Figure 4.**
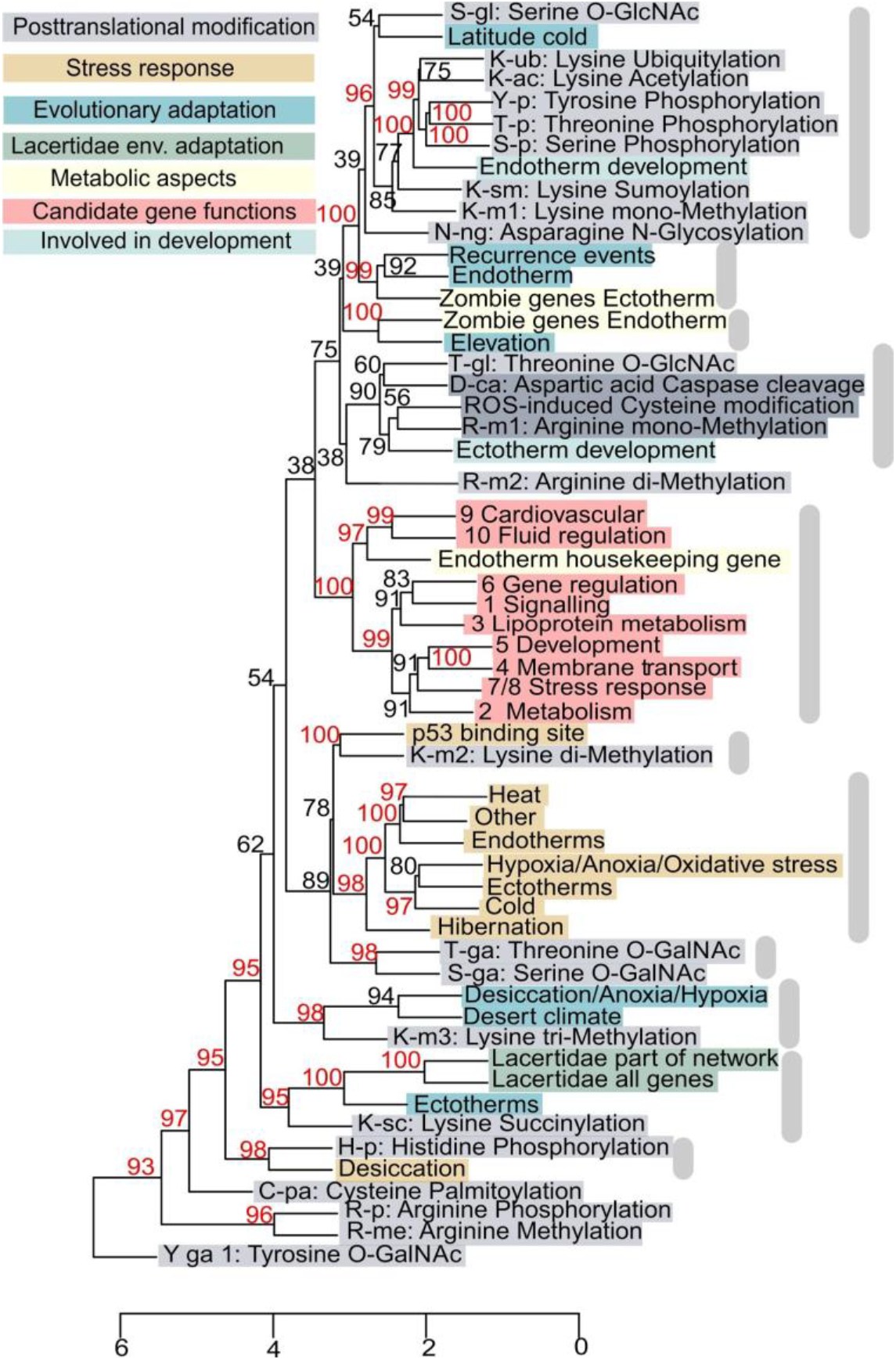
Results of Multiple Correspondence Analysis (MCA), represented as a dendrogram, to scale EAGs by different functional and evolutionary aspects. Statistical support was obtained via multiscale bootstrap (N=100,000 replicates (67)). 11 functional clusters were obtained with combined bootstrap support (alpha>0.9, red numbers on nodes), and standard error of estimate <0.01 (not shown).

## Discussion

In this contribution, we investigated the relative importance of genomic constraint vs contingency in shaping adaptation to the abiotic environment at the genome level. We expected that the existence of functional genomic constraint in adaptation could be proven through higher than random levels of identity and of functional similarity among genes that have adapted to similar selective regimes among different vertebrate taxa and physiological strategies. We here found evidence that this type of constraint plays an important role in generating environmental adaptation.

We found that EAGs of different species form a tight network with more functional connections among the genes than expected by chance. Across the different species and environmental factors considered, almost half of all genes in this functional network (43.6%) evolved recurrently, which was significantly higher than simulated in the absence of any constraint. Such events of recurrent evolution span a wide range of both ectotherm and endotherm species, as also a range of environmental factors. For example, 14 EAGs of Lacertidae that are related to a bioclimatic gradient have been independently recruited for environmental adaptation in other vertebrates. Ectotherms use these to adapt to desiccation and anoxia (killifish *Austrofundulus limnaeus)*, altitude (Himalayan *Nanorana* frogs, Himalayan *Phrynocephalus* lizards), and endotherms use them to adapt to latitude (Yakutian horse, woolly mammoth, minke whale, polar bear), deserts (camel and dromedary), and altitude (Himalayan marmot). Aspects of environmental adaptation were not clearly separated between processes in vertebrate endotherms and ectotherms, which indicates that the genomic mechanisms and functions of adaptation to abiotic parameters seem to be constrained even among these two, very different physiological strategies. This constitutes another indicator for the presence of genomic constraint in environmental adaptation. For example, Lacertidae shared in total five genes under selection with the extinct woolly mammoth (ETFA, MRPS31, PFAS, PHKB, and SENP5).

In lacertid lizards, terminal branches with very high values of episodic positive selection were identified mainly in species inhabiting cooler, more seasonal and partially montane environments such as *Podarcis muralis* and *Zootoca vivipara*. In contrast, they were absent in hot-arid adapted species such as *Mesalina olivieri* and species living in tropical environments (*Takydromus sexlineatus, Holaspis guentheri*). Adaptation in genes linked to mitochondrial cell respiration and oxidative stress response (proteasome component genes and the chaperone HSP90B1), could facilitate adaptation to cold environments via influencing the metabolic rate (32, 33). This corroborates previous findings that lower preferred temperatures are more likely to evolve than higher thermal preferences in lizards inhabiting climatic gradients (30, 34). Lacertidae genes under selection but not part of the vertebrate functional network did not have any functional enrichment, and could have evolved in response to selection pressures other than abiotic climate-related factors. Our comparative genomic approach thus may help disentangle environmental adaptation from covarying factors such as locomotor performance or reproduction (35).

The results with regards to gene function are in alignment with the hypothesis of constraint as well. High functional connectedness, and comparatively higher values of GO semantic similarity of the EAGs analyzed in our study indicated that they were drawn from a nonrandom set of very similar functions (36). Many EAGs fell within predicted candidate functions for climate adaptation (11, 31). Involvement in cytoskeletal and apoptotic processes, as well as localization and transport emerged as novel environmental adaptation-related function. Both apoptosis and transport-related genes have previously been associated with death-related processes, during which transport genes may become activated attempting to restore homeostasis (16). The cytoskeleton is remodelled through stress and involved in HSP phosphorylation (37) which in turn protect cytoskeletal integrity (38). Stress furthermore induces changes in cytoskeleton-regulated cell volume (39).

PTMs have previously been related to regulation of torpor as environment-induced metabolic process (40). Surprisingly, we found that specific PTMs were present in almost all functional clusters associated with environmental adaptation. Genes that were involved in latitudinal adaptation were related to various types of PTMs such as serine O-GlcNAc (serine-β-linked N-acetylglucosamine modification) which are associated with transcription, metabolism, apoptosis, organelle biogenesis, and transport (41) and disease including metabolic syndrome and cancer (41). 89% of all EAGs were involved in at least one human cancer, and a subset of them had cancer involvement as predominant function (e.g., FCGBP and SYT11 adapting to latitude in polar bear and woolly mammoth), which reinforces a relationship between abiotic environmental adaptation and homeostasis disturbance via disease (16). “Zombie genes” evolve recurrently, and are associated with altitude adaptation. These genes are both activated in dying cells, and involve localization and transport processes, which supports these functions as emerging cellular candidate processes for environmental adaptation. Adaptation to hypoxia and desiccation is furthermore associated with the PTM lysine trimethylation (42). Lastly, lysine succinylation was linked to genes involved in environmental adaptation in lacertids and other ectotherms, possibly related to UV radiation (43).

Our transcriptome data set on Lacertidae genes could be biased for genes with high expression (that are more likely recovered from all species in an RNA-based alignment), and further environmental adaptations could exist in other genes that we did not analyze; however, across all vertebrates, EAGs had higher expression values, higher numbers of PTM modification sites, and higher functional similarity than other partitions of the transcriptomes, but random sets of housekeeping genes had similar properties.EAGs have a higher expression level than expected by chance, which is similar to housekeeping genes, but are not identical with them. Only eight housekeeping genes were identified in our data set. A further 277 genes (or 25%) were, however, included in 3804 “wider definition” housekeeping genes, which are expressed at a more baseline or mid-level across all tissues and conditions (44). Genes with frequent expression in several tissues could expand the range of life stages and situations where natural selection can modify allele frequencies in response to abiotic environmental selection. However, as the dendrogram shows (Fig. 4), genes that are only expressed during embryonic development are aligned with thermal adaptation across latitude, elevation, and recurrent adaptation, pinpointing early development as a vulnerable stage for environmental selection (23).

Whilst 18% of environmental adaptation genes were involved in the response to stress in experimental studies, we did not find evidence for an alignment between specific axes of environmental adaptation to specific types of stress response such as desert adaptation to heat stress response. This result supports the idea that functional genetic networks are partitioned into different types of functions covered by distinct gene sets, which then interlink with others for information exchange (45), and supports the necessity of a generality of the organismal stress response to ensure viability in the face of multiple stressors (46). As expected, HSPs were differentially expressed under all stressors (47), and so were other genes functionally involved in the heat shock response. We identified little-studied EAGs, that ubiquitously responded to all analyzed stressors (PKLR, AHSG, SQRDL). Such genes may deserve further attention as possible biomarkers for environmental stress-related adaptation, as will be required under climate change conditions.

## Conclusions

Here we have shown that adaptations to abiotic environmental parameters are characterized by a high degree of genomic recurrence and functional similarity, in line with the hypothesis of genomic constraint. Although Lacertidae are thought to be excellent thermoregulators (48, 49), many lacertid lizard populations, especially those in humid montane localities, are currently undergoing population declines linked to climate change (50, 51). Studies are already underway that assess the future adaptive potential or “evolvability” of extant populations based on physiological parameters and distribution areas (52). The genes identified in this study deserve concerted focus in future research aimed at evaluating current climate stress on populations at the molecular level, and whether it may lead to evolutionary adaptation in the future. Bay and colleagues (53) pioneered such an approach to determine the likelihood of heat adaptation in corals. We recommend that similar efforts in vertebrates focus on the genes and functions outlined in this contribution, and on the specific functional mutations of different alleles within these candidate genes.

## Materials & Methods

Transcriptomic (RNAseq) data of 24 lacertid taxa were obtained from (29) (collection and ethics permits listed therein). The final alignment contained 6,269 gene sequences, which could be annotated to 5,498 unique gene symbols. Gene sequences obtained from RNAseq with no missing taxa were further checked for alignment errors including HMMCleaner for false negatives to pre-empt including artefacts in nucleotide-level analyses. Ten sequences were found to contain such potential artifacts and were removed from this data set. A maximum likelihood tree was generated from the concatenated alignment which is also published in (29). From this final dataset for 24 lacertids (22 species; two species with two divergent populations each) and one outgroup, we selected a subset of 695 genes with no missing sequences to analyze for signatures of selection. The aBSREL (54) random-effects branch site model as implemented in HyPhy (55) was used to find lineages subject to episodic positive diversifying selection. The full model was run in exploratory mode for all branches with Holm-Bonferroni correction. Summary statistics of dN/dS (ω) were computed across all branches where episodic positive diversifying selection was identified. A functional genetic network of all genes found to harbor signatures of selection was generated with Cytoscape v.3 (56) using the STRING app against the *Anolis carolinensis* genome as background, the closest related lizard with a sequenced and comprehensively annotated genome. The network was then tested for functional enrichment in Cytoscape using the ClueGO app v.2.5.0 (57), with the CluePedia plugin (58).

We compiled a matrix of putative EAGs in other vertebrate endotherms and ectotherms from 27 studies, adding the 200 genes obtained for Lacertidae. Support for this cross-taxon comparative functional genomics approach comes from the fact that orthologs across taxa are, despite their phylogenetic definition, most commonly identified by conserved function (59). The STRING database plugin within the Cytoscape (56) software was used to generate a functional genomic PPI network from these genes, using human PPI databases (59). We recorded and visualized whether single EAGs (nodes) within this network additionally adapted recurrently (i.e., in more than one species, whether they were responsive to stress, and recorded which of the Lacertidae genes under selection were part of this network. Gene functions in form of Gene Ontologies (GO), for EAGs that were additionally part of the resulting network (N=902 genes), were generated and grouped in ClueGo within Cytoscape (57). We then sorted, where appropriate, the resulting grouped GO terms into the functional categories for climate adaptation that we and others have previously defined (11, 12, 31). Additional GO functions were also recorded. We subsequently repeated this analysis for only those genes under selection in Lacertidae that were additionally part of the functional network (putative EAGs, N=171), and for those genes that were under selection in Lacertidae, but were not part of the functional network (less likely to be EAGs, N=29).

The 695 genes in our RNAseq alignment of Lacertidae represent a subset of genes of the entire aligned transcriptomes that were selected for analysis of selection based on the absence of paralogs and presence of transcripts in all 24 lacertid taxa, as explained above. Consequently, the 200 genes under selection found within this dataset might represent a nonrandom subset of the transcriptome dataset consisting of 5,498 annotated genes, with genes related to fundamental metabolic cell processes possibly being overrepresented as they would more likely to be found in all studied lacertid species. To test to which extent the functional properties of Lacertidae genes under selection is determined by a sampling bias and whether they are different from random genes, we therefore generated comparison groups in form of replicates of 200 randomly drawn genes. These were drawn from the total sequenced exomes (20 sets), from the genes not found under selection (20 sets), from 451 human housekeeping genes expressed in different types of healthy cells (brain, kidney, prostate, liver, muscle, lung, vulva, 10 sets) (60), and from the human genome (20 sets, obtained via www.molbiotools.com). GO terms are a hierarchical vocabulary-based classification method for gene functions (61). Functional similarity of gene products can therefore be quantitatively estimated through the semantic similarity of GO terms (61). We compared the mean semantic similarity of genes among the lacertid genes under selection, and the other comparison groups using ±95% prediction intervals with Z-tests. This analysis was repeated also for the semantic similarity of the 902 (lacertid + other vertebrate) EAGs that were part of the functional network, compared to 20 sets of 902 genes randomly drawn from the human genome as described above.

The question of constraint by gene identity was addressed by identifying the proportion of genes that were found to have adapted to differences in the abiotic environment in more than one species (=adapted recurrently). We simulated the number of genes that have to be drawn from a hypothetical genome to yield the same number of recurrent EAG, and tested for differences between observed number of genes and the simulated distribution using ± 95% prediction intervals and Z-test.

Additionally, data on candidate functions for environmental adaptation were collected for all EAGs. Genes were binary coded for their association with pre-defined candidate functions for environmental adaptation. Significance was based on FDR-corrected p-values of overrepresented GOs obtained with ClueGo. To code involvement in cellular pathways active in cancer as a candidate function for EAGs, gene expression data for 813 EAGs were obtained from microarray experiments of 89 cancers (62) and universal reference tissue. With respect to involvement in the candidate function apoptosis, “zombie genes” were identified in our data matrix (16). We binary-coded involvement in the acute response to stress caused by environmental factors from published information from 75-omics studies on the environmental stress response. Data on presence and type of experimentally confirmed PTMs was obtained from the PhosphoSite database (www.phosphosite.org, last accessed February 2019), and compared counts against the previously described 20 replicates of 200 randomly drawn genes from the human genome using Mann-Whitney U test and Z-statistic of randomized Mann-Whitney W statistic. To investigate differences between ectotherm and endotherm species, genes expressed during embryonic development for zebrafish as an ectotherm model species (63) and humans as an endotherm model species (64) were also binary coded. The resulting gene/ function matrix was subsequently analyzed via MCA in R (package FactoMineR (65) to align genes into functional clusters. Statistical support for MCA dimension groupings were obtained from 100,000 multiscale bootstrap replicates in the R package *pvclust (66).* For details see Supplementary Methods.

## Acknowledgements

The Deutsche Forschungsgemeinschaft (DFG) supported M.V. and J.M. (VE 247/11-1 / MU 1760/9-1). The University of Hull is thanked for funding the Molecular Stress in changing aquatic environments (MolStressH_2_O) research cluster (supporting K.W.V., P.B.A., and L.F.); other cluster members are thanked for helpful discussions. Hervé Philippe (Station d’Ecologie Expérimentale CNRS, Moulis, France) is thanked for help with sequence analysis. We acknowledge the Viper High Performance Computing facility of the University of Hull and its support team. Computations were partly carried out at the Swedish National Infrastructure for Computing (SNIC) at Uppsala Multidisciplinary Center for Advanced Computational Science (UPPMAX) under project SNIC 2017/7-275. S.S. acknowledges an Allam PhD studentship and A.B. acknowledges a PhD scholarship within the Endothelial cells in chronic disease research cluster at UoH. I.I. was supported by a postdoctoral Juan de la Cierva-Incorporación postdoctoral fellowship (IJCI-2016-29566) from Spanish Ministry of Economy (MINECO).

## Data deposition statement

Additional figures, tables and code is provided in Supplementary Materials. RNA-Seq data are available from the Sequence Read Archive (SRA Bioproject PRJNA543749). Binary data table for environmental adaptation genes and functions is provided as supplemental file in. csv format (Table S7).

## Author contributions

KWV, MV, JM, KBS designed research, KWV, JGP, II, SK performed research, OJG, PP, APT contributed new reagents or analytic tools, KWV, JGP, II, LF, AB, SFS, APT, PBA analyzed data, KWV and all co-authors wrote the paper

